# DNA from non-viable bacteria biases diversity estimates in the corals *Acropora loripes* and *Pocillopora acuta*

**DOI:** 10.1101/2023.11.16.567475

**Authors:** Ashley M. Dungan, Laura Geissler, Amanda Williams, Cecilie Ravn Gotze, Emily C. Flynn, Linda L. Blackall, Madeleine J. H. van Oppen

## Abstract

**Background:** Nucleic acid-based analytical methods have greatly expanded our understanding of global prokaryotic diversity, yet standard metabarcoding methods provide no information on the most fundamental physiological state of bacteria, viability. Scleractinian corals harbour a complex microbiome in which bacterial symbionts play critical roles in maintaining health and functioning of the holobiont. However, the coral holobiont contains both dead and living bacteria. The former can be the result of corals feeding on bacteria, rapid swings from hyper- to hypoxic conditions in the coral tissue, the presence of antimicrobial compounds in coral mucus, and an abundance of lytic bacteriophages.

**Results:** By combining propidium monoazide (PMA) treatment with high-throughput sequencing on six coral species (*Acropora loripes*, *A. millepora*, *A. kenti*, *Platygyra daedalea*, *Pocillopora acuta*, and *Porites lutea*) we were able to obtain information on bacterial communities with little noise from non-viable microbial DNA. Metabarcoding of the 16S rRNA gene showed significantly higher community evenness (85%) and species diversity (31%) in untreated compared with PMA-treated tissue for *A. loripes* only. While PMA-treated coral did not differ significantly from untreated samples in terms of observed number of ASVs, >30% of ASVs were identified in untreated samples only, suggesting that they originated from cell-free/non-viable DNA. Further, the bacterial community structure was significantly different between PMA-treated and untreated samples for *A. loripes* and *P. acuta* indicating that DNA from non-viable microbes can bias community composition data in coral species with low bacterial diversity.

**Conclusions:** Our study is highly relevant to microbiome studies on coral and other host organisms as it delivers a solution to excluding non-viable DNA in a complex community. These results provide novel insights into the dynamic nature of host-associated microbiomes and underline the importance of applying versatile tools in the analysis of metabarcoding or next-generation sequencing data sets.

## Background

Nucleic acid-based analytical methods, in particular 16S ribosomal RNA (rRNA) gene metabarcoding, have greatly expanded our understanding of global prokaryotic diversity. However, standard metabarcoding methods provide no information on the metabolic activity or physiological state of bacteria. Even the most fundamental physiological state, viability, cannot be assessed with conventional DNA-targeted methods such as PCR [1, 2].

One system with a complex microbiome is the coral holobiont [3], which includes the coral animal along with its diverse set of microbial symbionts [4, 5]. Of particular interest is the coral-associated bacterial community, which has been suggested to aid in the defence against pathogens [6, 7], nutrient cycling [8–13], the biosynthesis of essential amino acids [14, 15], and protection against climate change [16]. However, corals feeding on bacteria [17], rapid swings from hyper- to hypoxic conditions in the coral tissue [18], the presence of antimicrobial compounds in coral mucus [6], and an abundance of lytic bacteriophages [19–23] can contribute to a considerable prevalence of non-viable microbes and extracellular DNA (exDNA). exDNA and non-viable cells can persist in environmental samples for hours to months [24, 25]. This can bias or overwhelm analyses seeking to understand the diversity of coral-associated viable bacteria in the holobiont. While there are 295 studies using 16S rRNA gene metabarcoding to evaluate bacterial communities in coral samples (from Web of Science Core Collection search across all fields for: coral AND (metagenom* OR metabarcoding OR amplicon) AND bacteria), none have assessed bacterial community viability.

One method used for community viability characterization is high-throughput sequencing combined with a propidium monoazide (PMA) treatment to deplete signals from exDNA [26, 27]. PMA is not permeable to intact cell membranes; thus, the dye only interacts with DNA in membrane-compromised cells (*i.e.*, dead or damaged cells) or exDNA. Once the PMA is inside the cells it intercalates into the DNA and, after photoactivation, is crosslinked to the DNA, preventing PCR amplification [28]. PMA has been used in combination with PCR and 16S rRNA gene metabarcoding to selectively detect viable microbes without signals from damaged cells and exDNA in soil [26], sewage sludge [29], and guts of mammals [30–33] or fish [34, 35]. In these systems, DNA from non-viable microbes represented 9-40% of the amplified community and inflated community diversity by 25-55%. Overall, exDNA or DNA from non-viable microbes has been shown to obscure experimental treatment effects, spatiotemporal patterns, and relationships between microbial taxa and environmental conditions.

Here, we characterize the viable bacterial communities from six Great Barrier Reef (GBR)-sourced coral species, *Acropora loripes*, *A. millepora*, *A. kenti*, *Platygyra daedalea*, *Pocillopora acuta*, and *Porites lutea*.

## Methods

### Coral Sampling

All corals used in this study were part of the coral stock at the Australian Institute of Marine Science’s (AIMS) National Sea Simulator (SeaSim) in Townsville, Queensland, where they were housed in outdoor mesocosms with natural light conditions and a flow-through system with a full seawater exchange every two hours. Details about collection dates and locations are in Table S1. The daily temperature profile followed the average temperature profile recorded at Davies Reef weather station. Corals were maintained on a daily regimen of 0.5 *Artemia* nauplii ml^-1^ and 2000 cells ml^-1^ of a mixed-species microalgae solution. In June 2022, ∼2 cm fragments were cut from five different positions in each colony (distinct genotypes) of *A. loripes*, *P. daedalea*, *P. acuta*, and *P. lutea* without removing the colonies from the aquarium (n=25 each). Four genotypes of *A. kenti* and *A. millepora* were available in the SeaSim at the time of sampling (Table S1). GBR acroporids have recently undergone taxonomic revision; phylogenomic reconstructions consistently places ‘*A. tenuis*’ sampled from the GBR with other specimens from eastern Australia, identified as *A. kenti* [36]. Following this formal taxonomic revision, we have adopted this nomenclature. To minimise cross-contamination, nitrile gloves were discarded between each sampling, and all collection equipment (bone cutters and forceps) was sequentially sterilized in 10% sodium hypochlorite, reverse osmosis water, 80% ethanol, with a final wash in 0.22 µm filtered seawater (fSW) as described in Damjanovic et al. (2020).

Fragments were gently rinsed in fSW to remove loosely associated bacteria. Using an airbrush, tissue was removed from the fragment into a volume of 10 ml fSW, which was transferred to a 15 ml polypropylene tube. During the tissue airbrushing, 12 blanks were collected to account for contamination that might be introduced at this stage. Coral homogenate and blank samples were centrifuged (5,250x *g* for 10 min at room temperature (RT)), supernatant removed, and pelleted cells resuspended in 1000 µl 0.22 µm filter sterilized 3x concentrated phosphate buffered saline. Aliquots of 400 µl were transferred to two separate 1.5 ml tubes; the “A” tube to receive PMA treatment and the “B” tube as untreated. The untreated “B” tube was centrifuged (5,000x *g* for 10 min at RT), supernatant removed, and pelleted cells stored at −20 °C until shipment.

### PMA treatment

Extracellular DNA (exDNA) and DNA from membrane-compromised bacterial cells was removed by treatment of coral homogenates and tissue airbrushed blanks with the viability dye PMAxx^TM^ (Biotium, Fremont, CA, USA). PMAxx is a DNA-intercalating agent that forms photo-induced crosslinks making the bound DNA inaccessible for downstream molecular applications. PMAxx was added to the “A” fraction at a final concentration of 25 µM by adding 1 µl of 10 mM PMAxx stock to the 400 µl aliquot. This was followed by 10 min incubation in a rotary mixer inside a dark room at RT. Photoactivation was performed by using the PMA-Lite^TM^ LED Photolysis Device (Biotium) with the exposure time set to 30 min. During this exposure, samples were mixed every 3 min to ensure adequate photoactivation. After photoactivation, the PMA-treated “A” tube was centrifuged (5,000x *g* for 10 min at RT), supernatant removed, and pelleted cells stored at −20 °C until shipment.

The efficacy of PMAxx in cnidarian tissue was confirmed prior to coral collection at AIMS using a model for corals, the sea anemone *Exaiptasia* [37]. Triplicate heat-killed anemone samples (250 µl homogenized tissue heated to 90 °C for 60 min and plated to confirm no bacterial growth) were spiked with either 3.7 × 10^6^ heat-killed or viable *Endozoicomonas* sp. cells (strain ALC013, provided by Cecilie Goetze). Additional triplicate heat-killed anemone samples were included without addition of bacteria. Subsequently, samples (n=12) were split into two aliquots of which one was treated with PMAxx as described above and one left untreated. DNA was extracted from these samples (as below) and bacterial load quantified using droplet digital PCR (ddPCR) targeting the bacterial 16S rRNA genes. DNA extraction and library prep PMA-treated and untreated samples (n=302) were shipped to The University of Melbourne for DNA extractions, which were completed as described by Hartman LM, van Oppen MJH and Blackall LL [38]. Blank DNA extractions (n=9) were included to identify reagent contamination.

Extracted DNA was amplified using bacterial primers targeting the V5-V6 regions of the 16S rRNA gene: 784F [5′ <u>GTGACCTATGAACTCAGGAGTC</u>AGGATTAGATACCCTGGTA 3′], 1061R [5′ <u>CTGAGACTTGCACATCGCAGC-</u>CRRCACGAGCTGACGAC 3′] (Andersson et al 2008) with overhang adapters (underlined). Negative template PCR controls (n=7) were included to test for potential contamination. Triplicate PCRs were carried out in 15 µl reactions containing 1x UCP Multiplex PCR Master Mix (Qiagen, Venlo, The Netherlands), 0.3 μmol l^−1^ each of the forward and reverse primers, and 1 µl of template DNA (1:40 dilution) or nuclease-free water (negative template control). The PCR cycling conditions consisted of an initial heating step at 95 °C for 3 min; 18 cycles of 95 °C for 15 s, 55 °C for 30 s and 72 °C for 30 s; and a final extension step of 72 °C for 7 min. Triplicate PCR products were then pooled; successful DNA extraction was confirmed by agarose gel electrophoresis.

A volume of 20 µl of each PCR product pool was purified by size-selection using Nucleomag NGS Clean-up and Size Select beads (Scientifix, Clayton, VIC, Australia). The purified DNA was resuspended in 40 µl of nuclease-free water. Indexing PCRs were created by combining 10 μl of purified DNA with 10 μl 2x MyTaq HS Mix polymerase (Bioline, Narellan, NSW, Australia) and 1 μl (5 μM) of forward and reverse indexing primers. The PCR cycling conditions consisted of an initial heating step at 95 °C for 3 min; 24 cycles of 95 °C for 15 s, 60 °C for 30 s and 72 °C for 30 s; and a final extension step of 72 °C for 7 min. For a subset of randomly chosen samples, product size was confirmed by agarose gel electrophoresis. Barcoded PCR amplicons were pooled by species; pools were purified with a bead clean up. Each pool was checked for quality and quantity (2200 TapeStation; Agilent Technologies, Mulgrave, VIC, Australia) to guide pool normalization at equimolar concentrations, then sequenced on a single Illumina MiSeq run using v3 (2 × 300 bp) reagents at the Walter and Eliza Hall Institute, Melbourne, Australia.

### Metabarcoding

Raw demultiplexed 16S rRNA gene sequences were imported into QIIME2 v2022.11 [39] where primers and overhang adapters were removed [using cutadapt v2.6; 40]. The data were then filtered, denoised using the pseudo-pooling method, and chimera checked [using DADA2 v1.22.0; 41] to generate amplicon sequence variants (ASVs). Taxonomy for each ASV was assigned against a SILVA database (version 138) trained with a naïve Bayes classifier against the same V5-V6 region targeted for sequencing [42]. A phylogenetic tree was produced in QIIME2 by aligning ASVs using the PyNAST method [43] with mid-point rooting.

### Droplet digital PCR

ddPCR was used to track changes in the absolute abundance of bacteria and the *Endozoicomonas* community between PMA treated and untreated samples for *A. loripes* only. ddPCRs for each sample (n=50) were prepared in an initial volume of 72 µl comprising 37.5 µl EvaGreen Supermix (QX200; Bio-Rad, South Granville, NSW, Australia), 31.5 µl sterile water, and 3 µl diluted DNA template. The solution was mixed and then split into three 24 µl aliquots, one for host cell quantification (*A. loripes* β-actin gene), one for all bacteria, and a third to target *Endozoicomonas* spp. The β-actin gene occurs as a single copy in *A. loripes* and is used here as an internal control gene. Primers to target the *A. loripes* β-actin gene were designed using Primer3 [v4.1.0, 44] using the β-actin gene sequence from the existing draft genome [45]. One microlitre each of the appropriate 5 µM forward and reverse primers (Table 1) were added to each reaction aliquot, giving final primer concentrations of 200 nM and volumes of 25 µl. From each 25 µl volume, 20 µl was loaded into a DG8 cartridge (1864008; Bio-Rad), followed by 70 µl of droplet generation oil for EvaGreen (1864005; Bio-Rad), and droplets were generated in a droplet-generator (QX200; Bio-Rad). A volume of 40 µl of generated droplets per reaction was then transferred to a 96-well plate and foil-sealed (1814040; Bio-Rad) with a thermal plate sealer (PX1; Bio-Rad). No-template controls and duplicate samples were included in each plate. Thermal cycler settings were: one cycle at 95.0 °C for 6 min; 40 cycles at 95 °C for 15 s + 56 °C for 15 s + 72 °C for 15 s; one cycle at 72 °C for 4 min; 12 °C hold.

**Table 1:**
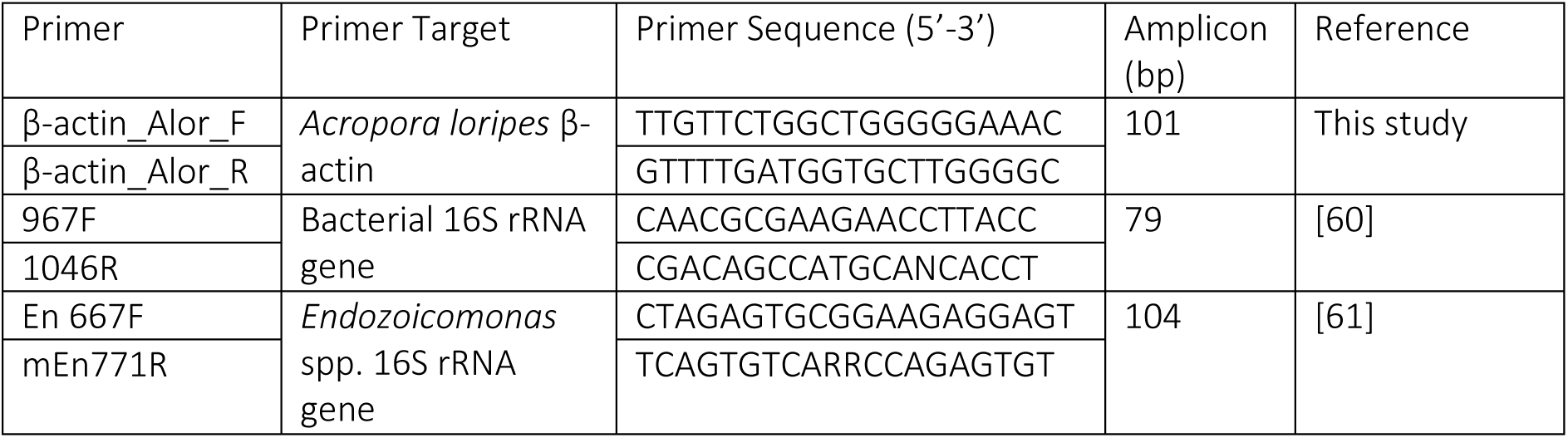
Primer sets used for ddPCR.

Droplets were read on a Bio-Rad QX200 droplet reader, and fluorescence data were analysed in QuantaSoft v1.7.4.0917. To quantify the change in *Endozoicomonas* abundance in coral cells, the 16S rRNA gene of *Endozoicomonas* was measured using *Endozoicomonas*-specific primers and normalized using the coral β-actin gene. To quantify the variation in total bacterial abundance in coral, the 16S rRNA gene in bacteria was quantified using bacterial primers and normalized with the β-actin gene. To determine the dynamics in relative abundance of *Endozoicomonas* to the total bacterial community, the *Endozoicomonas* 16S rRNA gene was measured using *Endozoicomonas*-specific primers and normalized by the 16S rRNA gene of total bacteria.

### Statistical analysis

All data were analyzed in R [v4.2.3; 46]. For metabarcoding data, ASV, taxonomy, metadata and phylogenetic tree files were imported into R and combined into a phyloseq object [47]. Contaminant ASVs were identified and removed sequentially from the dataset according to their abundance in the PCR negative controls, extraction blanks, and tissue processing blanks relative to the samples using the prevalence method in the R package decontam with p=0.1 [48].

α-diversity metrics (observed ASVs, Shannon’s and inverse Simpson’s indices) were calculated from a rarefied dataset by species using the ‘estimate_richness’ function in the R package phyloseq [47]. These and the ddPCR data were then analyzed using linear mixed effects models, with PMA treatment and coral genotype as fixed effects, and aquarium as a random effect, using the R package nlme [49]. Where appropriate, *post hoc* comparisons were performed using Tukey’s honestly significant difference test in the R package emmeans [50]. Box plots were created with ggplot2 [51], plotrix [52], and gridExtra [53] by merging data by PMA treatment. Summary data was produced using the function ‘summarySE’ from the package Rmisc [54].

β-diversity was evaluated using a weighted Unifrac distance matrix. The difference in community compositions among groups (PMA treatment and coral genotype) was calculated using ‘adonis2’ (a modified version of a permutational multivariate analysis of variance (PERMANOVA)) in the vegan package in R [55] with ‘Tank’ assigned as a random effect and 9999 permutations. Where the assumption of homogeneity of dispersion was not met, a permutation test for multivariate dispersion to account for unequal dispersion between groups was included. Where appropriate, Holm corrected pairwise comparisons were computed using the package pairwiseAdonis [56].This method tests the hypothesis that the bacterial communities from different groups (e.g., PMA treatment and genotype) are significantly different from each other in terms of their composition. PERMANOVA uses a distance matrix as input and is often more robust to non-normality and heteroscedasticity than traditional ANOVA. The results of the PERMANOVA were cross validated with and visualized using principal coordinates analysis (PCoA) to gain a more robust understanding of the underlying patterns. Where there were significant differences in community composition between PMA treatments for a given species, ASVs that were significantly associated with treatment groups were identified using the indicspecies function ‘multipatt’ [57] with specificity (At) and fidelity (Bt) set to 0.6. Barplots visualizing overall microbiota composition were made using ggplot2 [51] by agglomerating taxa at the genus or ASV level based on relative abundances. Venn diagrams were created using packages eulerr [58] and microbiome [59] to identify ASVs in PMA-treated or untreated samples for all corals and for each species.

For all tests p-values <0.05 were considered significant.

## Results

### Method validation

Prior to collecting coral samples, we examined the efficacy of PMAxx treatment in removing DNA from non-viable bacteria in a complex cnidarian matrix. We used heat-killed anemone tissue as our model system to mimic the complexity and opacity of homogenized coral tissue, while removing the potential influence of a living bacterial community. *Endozoicomonas* sp. was used to supplement the tissue and provide signal as it is one of the most ubiquitous bacterial symbionts of corals. Spiking of heat-killed anemone samples with heat-killed or live *Endozoicomonas* showed that PMAxx treatment effectively eliminated DNA from non-viable bacteria; there was a substantial reduction in copies of the 16S rRNA gene (per µl) in ddPCR (using general bacteria probes) when PMA was applied to anemone tissue spiked with heat-killed bacteria (mean±SE, 68.2±26.3) compared to the untreated fraction (2358.7±225.8), equivalent to a signal reduction of 97.1%. This copy number was largely recovered when anemone tissue was spiked with live *Endozoicomonas* in parallel with PMA treatment (Fig. S1), though signal after bacterial addition with PMA treatment was only half that of the untreated fraction suggesting some bacteria in the added culture may have not been viable.

### Metabarcoding data

Sequencing produced 10.7 M reads between the six coral species (n= 290), sampling blanks (n=12), extraction blanks (n=9), and PCR negative control samples (n=7). After merging, denoising and chimera filtering, 6.8 M reads remained. ASVs with fewer than 10 reads were removed from the dataset as they likely represent sequencing errors. Decontam identified eight putative contaminant ASVs from PCR amplification (0.02% of total reads), eight putative contaminant ASVs from DNA extraction (0.07% of total reads), and 11 putative contaminant ASVs from tissue blasting blanks (0.50% of total reads) (Table S2). Nine of 290 samples (four *A. loripes*, one *A. millepora,* four *P. daedalea*) were removed from analysis as they had <500 reads. After all filtering steps, there were 8,499 ASVs across the remaining coral samples (n=281).

Coral species were subset into separate phyloseq objects for analysis, and independently rarefied (Table 2) based on rarefaction curves for species richness (Fig. S2). PMA treatment had a significant effect on α-diversity indices for the microbiota present in *A. loripes* only. The evenness (inverse Simpson’s index) in untreated (mean±SE) (9.20±1.12) compared with PMA treated (4.97±0.55) samples showed an 85% inflation as result of non-viable DNA. Likewise, Shannon’s Index of diversity (2.49±0.10) in untreated samples was overestimated by 31% compared to the PMA treatment (1.90±0.12) (Fig. 1; Table 2).

**Figure 1:**
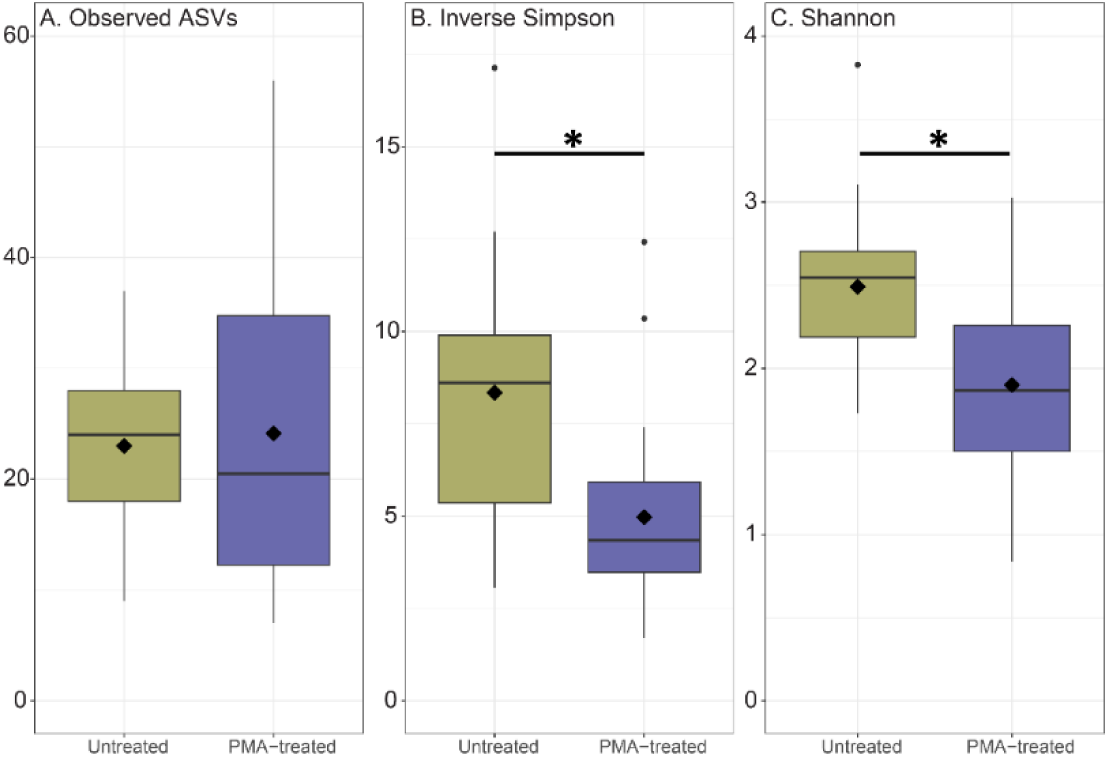
α-diversity indices for *A. loripes*, A) observed ASVs B) inverse Simpson’s index and C) Shannon’s index. Boxes cover the interquartile range (IQR) and the diamond inside the box denotes the median. Whiskers represent the lowest and highest values within 1.5 × IQR. * indicates a significant difference between treatments.

**Table 2:**
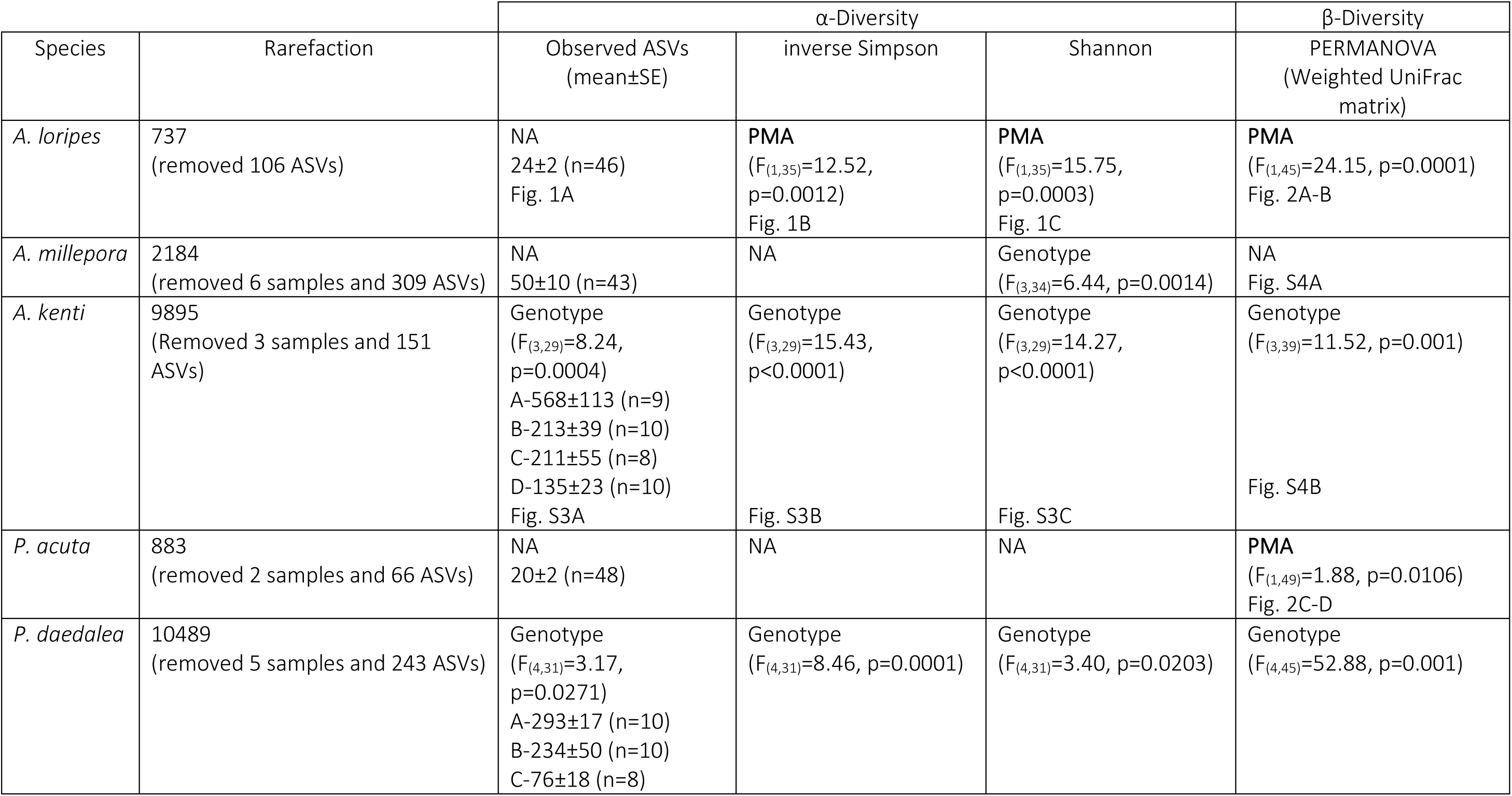
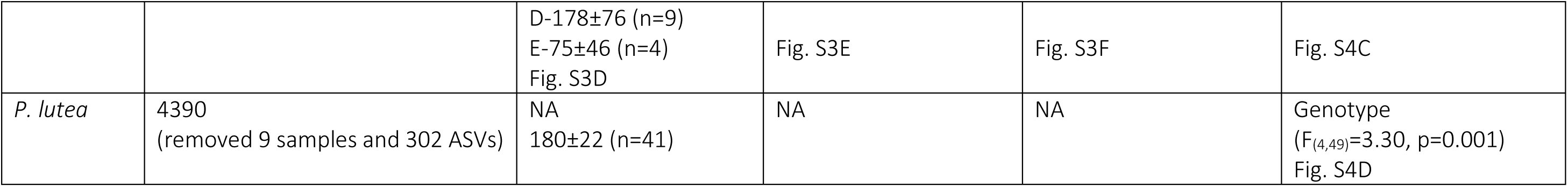
α- and β-diversity analysis results by coral species. Rarefaction levels were chosen based on rarefaction curves for Observed ASVs (Fig. S2). A significant main factor effect is denoted by “PMA” or “Genotype;” there were no interactions between main effects. Where there were significant differences, statistics are provided as well as the figure where a given data type is visualized.

For *A. kenti* and *P. daedalea*, there was a significant effect of host genotype on all α-diversity parameters (Table 2; Fig. S3), with one genotype largely driving the differences (A for *A. kenti* and B for *P. daedalea*). Only Shannon’s index of diversity was significantly different by genotype for bacterial communities in *A. millepora* (Table 2). While PMA-treated corals did not differ significantly from untreated samples in terms of mean ASV richness for any species, of the 8,499 ASVs in the full coral dataset, >30% were identified in untreated samples only, alluding that they originated from exDNA or non-viable DNA (Fig. 2A).

**Figure 2:**
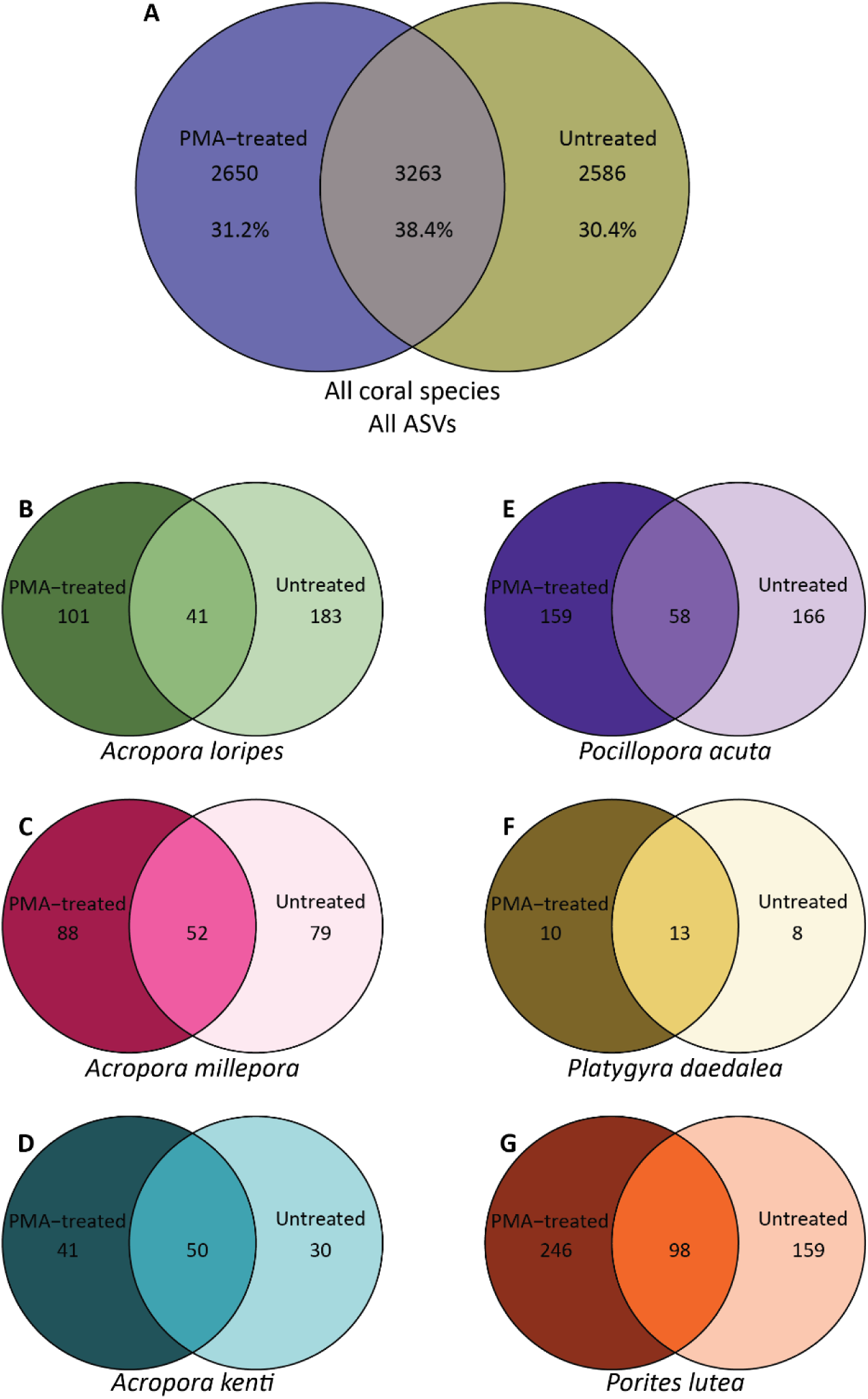
Venn diagrams showing the total number of ASVs in PMA-treated (left), untreated (right), or shared (centre) for all corals (A), *A. loripes* (B), *A. millepora* (C), *A. kenti* (D), *P. acuta* (E), *P. daedalea* (F), and *P. lutea* (G). For all corals, this includes all ASVs in all coral samples. For each species-specific diagram, only ASVs that made up at least 0.1% of the microbiome in a given sample were included in the count.

PERMANOVA analysis revealed a significant effect of PMA treatment on bacterial community structure for *A. loripes* and *P. acuta* samples, regardless of genotype (Table 2), which is illustrated by the distribution of samples in the PCoA of weighted unifrac distances and community composition (Fig. 3). The communities of both PMA treated and untreated *A. loripes* were dominated by *Endozoicomonas* spp.; however, the relative abundance of *Endozoicomonas* shifted from 29.5% in the untreated to 78.2% in the PMA treated samples (Fig. 3A). Untreated *A. loripes* had 24-fold greater relative abundance of an unknown Rickettsiales (26.4% untreated, 1.1% PMA-treated) and 5x more *Candidatus* Fritschea, a member of the *Simkaniaceae* family (3.7% untreated, 0.7% PMA-treated). PMA treated samples had 7-fold more *Vibrio* (0.4% untreated, 3.0% PMA-treated) and saw the emergence of *Acinetobacter* (0.002% untreated, 4.0% PMA-treated). For *P. acuta* (Fig. 3C), PMA treatment resulted in a loss of *Reinekea* (15.7% untreated, 0% PMA-treated) and *Nonlabens* (8.2% untreated, 2.0% PMA-treated), with increases in *Salinipshaera* (1.8% untreated, 8.8% PMA-treated), an unknown *Oceanospirillales* (2.3% untreated, 5.4% PMA-treated) and the appearance of XBB1006 (0% untreated, 6.1% PMA-treated). The bacterial β-diversity of *A. millepora*, *A. kenti*, *P. daedalea*, and *P. lutea* was not impacted by PMA treatment, though there were differences in the presence of individual ASVs by PMA treatment (Fig. 2) and all but *A. millepora* had a significant impact of host genotype on bacterial community composition (Table 2, Fig. S4. Table S3).

**Figure 3:**
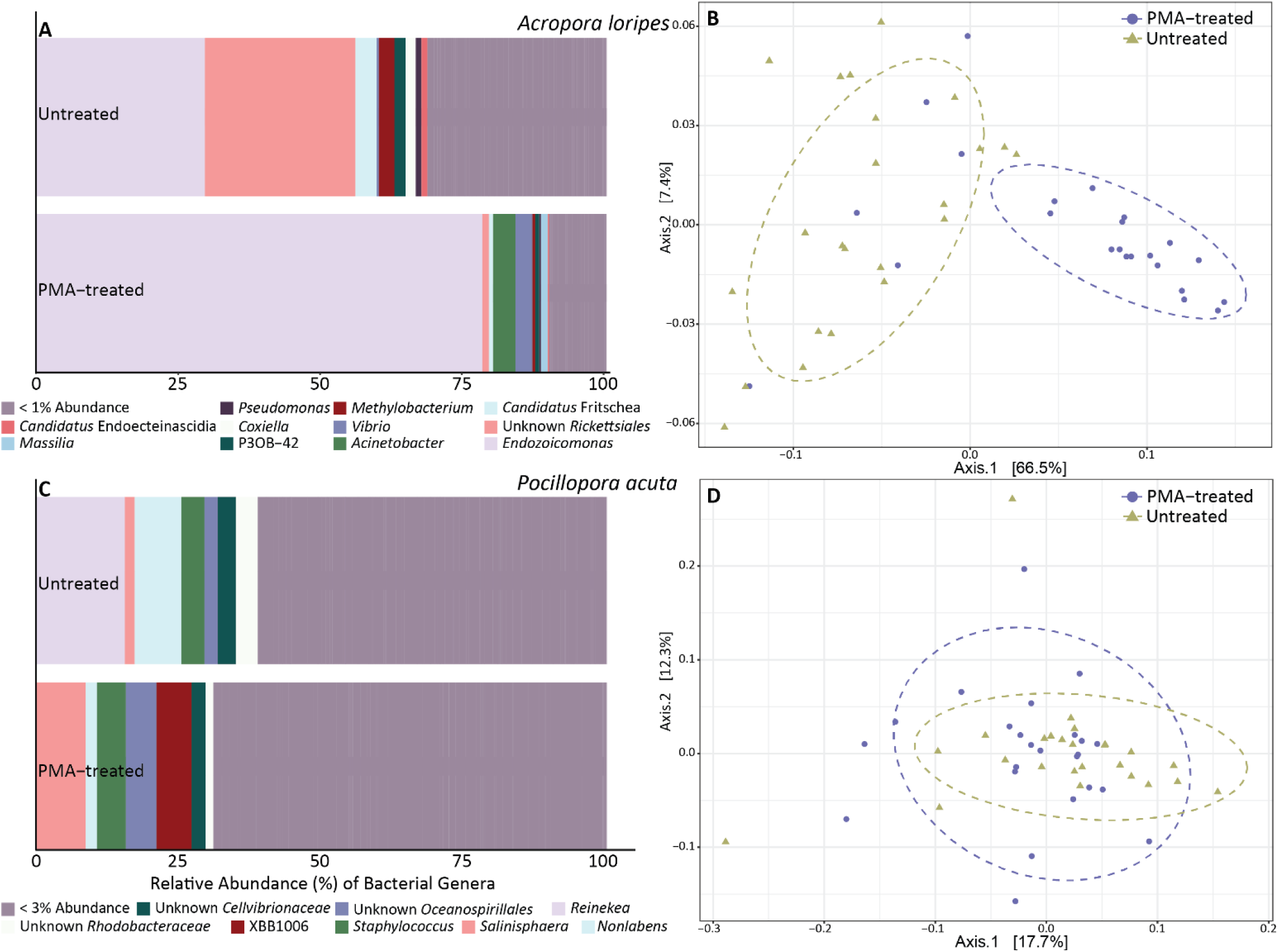
Relative abundance of bacterial genera for *A. loripes* (A) and *P. acuta* (C) samples in the Untreated and PMA-treated groups. Genera whose relative abundance was less than 1% or 3%, respectively, in both treatments were pooled into a single category. Principle coordinate analysis (PCoA) for *A. loripes* (B) or *P. acuta* (D) tissue using a weighted UniFrac distance matrix for untreated (gold triangles) and PMA-treated (purple circles) samples with ellipses drawn at 95% confidence levels for a multivariate t-distribution.

The indicator species analysis provided two ASVs characteristic of the microbiomes of the PMA-treated or untreated samples for *A. loripes* and one ASV for *P. acuta* (Table 3). ASV001 was identified as an indicator for the PMA-treatment in *A. loripes*; BLASTn output for this sequence provided *Endozocomonas coralli* and *E. atrinae* as top hits at 99.22% identity. There was a 2.3-fold increase in ASV001 from the untreated to the PMA-treated condition. This ASV was also found, albeit in low relative abundance, in *A. millepora* (Fig. S5A-B). ASV002 is an unknown member of the class *Alphaproteobacteria*, potentially belonging to the order *Rickettsiales* or *Rhodospirialles*. This ASV was an indicator of the untreated condition in *A. loripes* where it decreased 23-fold when samples were PMA treated (Table 3). Only one ASV was identified as an indicator taxon for *P. acuta*, ASV003. This ASV was a 100% match to *Reinekea thalattae* and, while it made up 15.1% of the abundance of all bacteria in the untreated condition, it virtually disappeared with PMA-treatment (0.002% relative abundance).

**Table 3:**
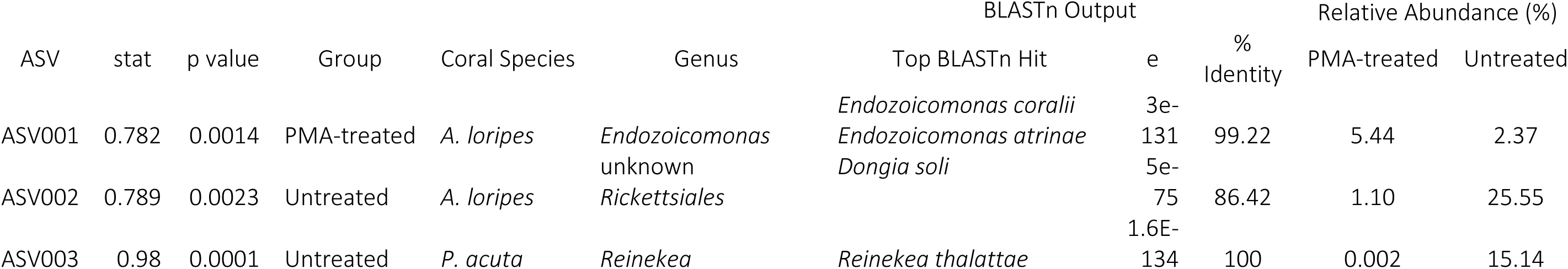
ASVs identified as being characteristic of either the PMA or untreated (Group) samples for *A. loripes* or *P. acuta* using an indicator species analysis. For the identified ASVs, the top BLASTn hit is reported with e-value and % identity for the match. The relative abundance (%) of the ASV is reported for each group in the respective coral species.

*A. loripes* had 72 *Endozoicomonas* ASVs between all the samples (Fig. S5A); ASV001 was the only variant identified as being indicative of PMA treatment. Other coral species were also dominated by *Endozoicomonas* including *A. millepora* (70 ASVs), *A. kenti* (33 ASVs), and *P. daedalea* (33 ASVs) (Fig. S5). Four of the *P. daedalea* genotypes were dominated by one ASV (ASV023; Fig. S5D), an unknown *Rhodanobacteraceae* who’s top BLASTn hit (98.07% identity) was *Fulvimonas yonginensis*.

### ddPCR

Given the significant effect of PMA-treatment on α- and β-diversity for bacterial communities associated with *A. loripes*, particularly *Endozoicomonas* spp., these samples were further processed to quantify bacterial load. For each sample, three data points were collected using ddPCR: 1) bacterial 16S rRNA gene copies per coral cell; 2) *Endozoicomonas* 16S rRNA gene copies per coral cell; and 3) the relative abundance of *Endozoicomonas*. Contrary to expectations, PMA treated samples had significantly more bacterial gene copies (F_(1,36)_=38.74, p<0.0001) and *Endozoicomonas* (F_(1,37)_=15.74, p=0.0003) per coral cell, but there was no difference in the ratio of *Endozoicomonas* to bacterial signal between the two groups (Fig. 4). There was no effect of *A. loripes* genotype on any of the ddPCR data.

**Figure 4:**
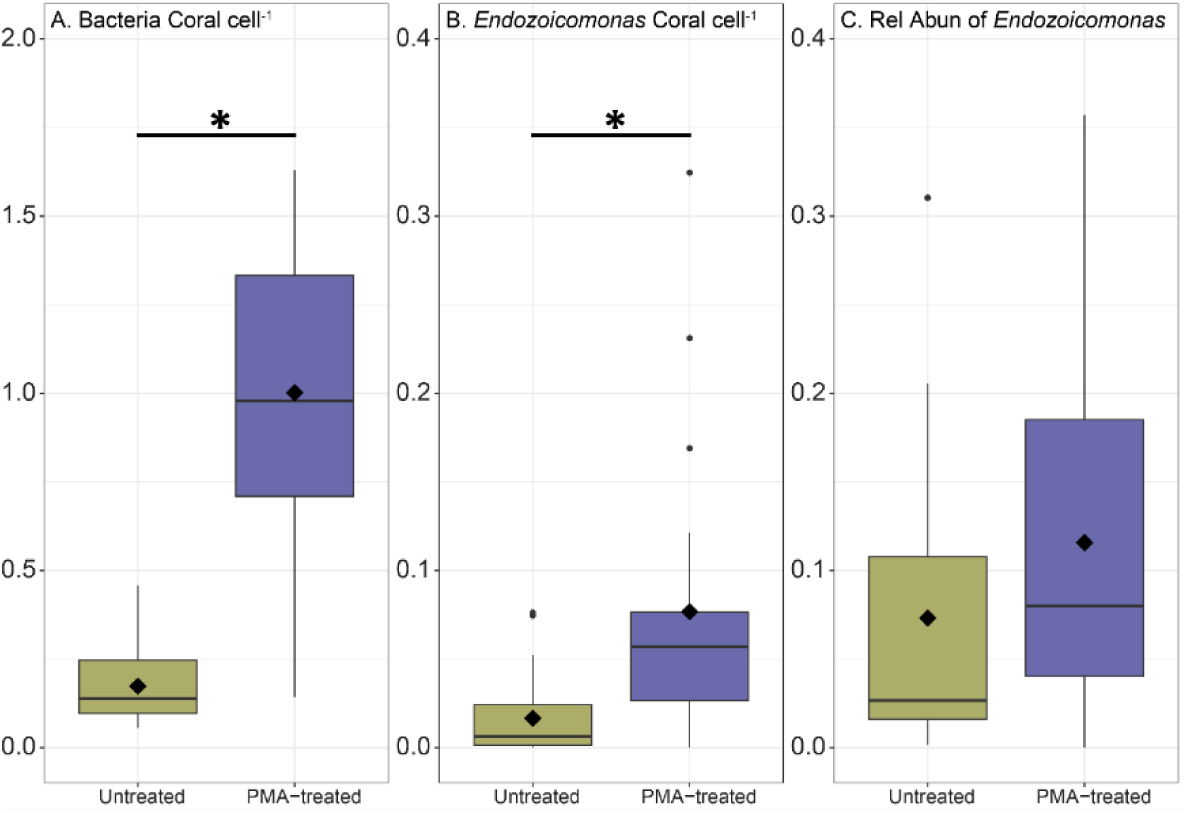
Bacterial load quantified using ddPCR showing, A) bacteria coral.cell^-1^ b) *Endozoicomonas* coral.cell^-1^ and C) relative abundance of *Endozoicomonas* (ratio of *Endozoicomonas* to total bacterial 16S rRNA gene copies), where 0.1=10%. Boxes cover the interquartile range (IQR) and the diamond inside the box denotes the median. Whiskers represent the lowest and highest values within 1.5 × IQR. * indicates a significant difference between treatments.

## Discussion

### Viability testing shows significant differences in bacteria associated with low diversity corals

Metabarcoding is often non- or semi-quantitative. DNA extraction and amplification bias, 16S rRNA gene copy number variation, bioinformatics classification accuracy, and (non)viability all influence the degree to which metabarcoding truly quantifies a community’s taxonomic profile. The use of 16S rRNA/16S rRNA gene ratio to identify viable bacteria in the coral system is gradually appearing [16, 62]. However, this approach shows inconsistent results in other systems [63]. Though not interpreted as such, Sun F, Yang H, Wang G and Shi Q [64] indirectly looked at the variation in coral-associated bacterial communities based on viability when comparing metatranscriptome and metagenome data, finding significant differences in the microbial community composition at DNA and RNA levels. This study is the first to apply PMA in combination with metabarcoding to coral tissue samples and shows that DNA from non-viable microbes significantly inflates community evenness (85%) and species diversity (31%) in *A. loripes* and bacterial community composition for *A. loripes* and *P. acuta*. This exaggeration in diversity with the presence of non-viable bacteria has previously been reported in a range of systems from rabbits to rainwater (Table S4), but for the first time here for corals.

PMA significantly altered α- and β-bacterial diversity metrics in *A. loripes* and β-diversity in *P. acuta*. Average bacterial richness in adult corals can range from 20 to >500 ASVs [65, 66]. *A. loripes* and *P. acuta* are on the lower end of this spectrum with (mean±SE) 24±2 and 20±2 ASVs per sample, respectively. The variation in response to PMA treatment by coral species could be explained by differential and specific antibacterial capabilities of the surface mucus layer [67], whereby some species may be more proficient at degrading environmentally sourced bacteria. Alternatively, coral species may have unequal expression of immune pathway genes, such as those involved in the Toll-like and nucleotide-binding oligomerization domain-like receptor signalling pathways, resulting in a range of responses to bacterial exposure. Low diversity corals may have a more active role in selecting their bacterial symbionts and have immune mechanisms in place to influence community composition. Further, since we did not quantify the number of bacteria per coral cell for all species, it is possible that low diversity corals had fewer bacteria all together and that eliminating exDNA and DNA from dead bacteria had a greater influence on the abundance of other viable members. These results correspond with previous studies that show bacterial communities post-PMA treatment had significantly reduced α-[29, 31, 33, 34] or β-[31, 34, 68] diversity indices, suggesting that standard 16S rRNA gene metabarcoding methods miscalculate coral-associated bacterial diversity for low diversity systems. For *A. loripes* the PCoA (Fig. 3B) shows greater variability in community composition for untreated compared to PMA-treated corals. This suggests that DNA from non-viable microbes may obscure effects of treatments between bacteria and environmental conditions for this species by exaggerating diversity.

### The differences between untreated and PMA treated bacterial communities were driven by a few taxa

For *A. loripes*, the reduction in diversity and evenness of bacterial communities was largely reflected in the abundance of *Endozoicomonas* spp. As non-viable bacteria and exDNA signal were removed, *Endozoicomonas* increased from 29.5% to 78.2% relative abundance in the untreated to PMA-treated condition. This suggests that *in hospite Endozoicomonas* are alive with few dead cells. Correspondingly, indicator species analysis identified ASV001, an *Endozoicomonas* sp., indicative of PMA treatment. In addition to ASV001, there were 71 other *Endozoicomonas* ASVs in *A. loripes* (Fig. S5A). The dominance of *Endozoicomonas* spp. in the viable bacterial community of *A. loripes* adds a valuable component to other work showing *Endozoicomonas* are a key symbiotic partner in this species [69, 70], other acroporids [71–76], and a global distribution of corals [11, 77–80].

Bacterial viability studies have found that previously identified “key” bacterial taxa were not active in the community sampled (Legrand et al. 2021). Here were saw a depletion of *Rickettsiales* in the viable bacterial community of *A. loripes*, a member of which was identified as an indicator taxon for the untreated samples (ASV002). *Rickettsiales* are consistently found in *Acropora* spp. [81–85] and corals globally [86]. This taxon is thought to be associated with white band disease [87] with genomic signatures of pathogenicity [86], though Casas V, Kline DI, Wegley L, Yu Y, Breitbart M and Rohwer F [81] found *Rickettsiales* to be widespread in both healthy and diseased *A. cervicornis* and *A. palmata* in the Caribbean. The absence of *Rickettsiales* in the viable bacterial community of *A. loripes* in contrast to their abundance in the untreated condition implies that they were predominately dead cells or exDNA. Their abundance could be explained by changes in environmental conditions that allow *Rickettsiales* to proliferate, coral immune responses, acquisition of *Rickettsiales* exDNA from the external environment, or feeding. Coral-associated *Rickettsiales* thrive under high nutrient conditions [88]; *Rickettsiales* could have bloomed under previous conditions and crashed in low nutrient load tanks. Alternatively, it is possible that acroporids have developed strategies to prevent infection by *Rickettsiales* leading to an abundance of dead cells. Members of *Rickettsiales* have been identified as infecting coral hosts through horizontal transmission [89], suggesting that corals may acquire this taxon from the surrounding seawater, possibly though heterotrophic feeding. Corals are known to heterotrophically feed on available bacteria [90–92] and *Rickettsiales* are found in tropical marine waters surrounding coral reefs [93].

ASV003 (*Reinekea thalattae*) was identified as an indicator taxon for untreated *P. acuta*, where it made up 15.1% of the microbiome and was depleted with PMA-treatment (0.002% relative abundance). *Reinekea* spp. have been previously reported in the coral-associated microbiome [94], potentially associated with disease [95], but there is otherwise little information about this taxon and its relationship to the coral holobiont.

### Not all coral species harbour a distinct viable microbial community, but there are differences

Four of the six coral species tested did not have significant differences by PMA treatment for bacterial α- or β-diversity. However, there were clear differences in the ASVs found in PMA-treated versus untreated samples, where 24.8% (*A. kenti*) – 36.1% (*A. millepora*) of the unique reads were found only in untreated samples. Since this value considers all ASVs in all samples (Fig. 2A) or all ASVs for each coral species (Fig. 2B-G), as opposed to the average number of ASV’s per sample (α-diversity richness measure), this suggests that, particularly for the rare/low abundant taxa, the ASVs are different between PMA-treated and untreated fractions. One hypothesis is that specific more highly abundant bacterial sequences are preferentially amplified in the untreated samples. A reduction in the abundance of these sequences following PMA treatment would then allow primers to bind to low abundant templates that were previously unbound. This is in line with observations that PMA treatment increases evenness and diversity without a change in the number of ASVs, a phenomenon driven by increased detection of rare taxa [30, 31, 96]. We also found that that coral microbiota varied by genotype for *A. kenti*, *P. daedalea*, and *P.* lutea, which has previously been reported [76, 85, 97].

Coral tissue-associated bacteria can form dense clusters termed cell-associated microbial aggregates (CAMAs). CAMAs have previously been visualized in all coral genera used in this study - *Acropora*, *Platygyra*, *Pocillopora*, and *Porites* [77, 78, 98–107]. One consistent trend is that *Endozoicomas,* a widespread coral symbiont [108], reside in CAMAs. No studies have yet visualized CAMAs in *A. kenti*, but the consistency and abundance of *Endozoicomonas* in this study and others suggest that CAMAs can form in this species. CAMA formation in *P. daedalea* has not been described since Work TM and Aeby GS [102] but the high abundance (>75%) of ASV023 in *P. daedalea* (Fig. S5D) is consistent with the presence of CAMAs. ASV023 is an unknown *Rhodanobacteraceae* who’s top BLASTn hit (98.07% identity) was *Fulvimonas yonginensis*. This family has been found in the mucus of *A. millepora* [109], but otherwise little literature exists on its relationship with coral.

### Eukaryotic DNA influences ddPCR results and can skew normalized bacterial results

ddPCR, qPCR, or fluorescence activated cell sorting (FACS), are elegant approaches for quantitative microbiome profiling, which can complement metabarcoding findings and can be used to track the amount of microbial DNA derived from non-viable bacteria. However, it is important to realize that technical sources of variability may introduce substantial additional biases depending on the quantification method used [33]. As with qPCR, ddPCR uses Taq polymerase in a standard PCR to amplify the target DNA. The ddPCR technology, however, partitions the PCR into thousands of droplets (individual reaction vessels) prior to amplification and acquires the data at the reaction end point. This enables more precise and reproducible data and direct quantification without the need for standard curves (Kim et al., 2014; Gobert et al., 2018). ddPCR results with *A. loripes* show that the number of bacteria per coral cell and *Endozoicomonas* per coral cell were significantly greater in the PMA-treated samples, with no difference in relative abundance of *Endozoicomonas*. This finding is surprising because the metabarcoding data showed an increase in *Endozoicomonas* relative abundance following PMA treatment. Given that PMA treatment blocks non-viable bacterial DNA from being amplified, the opposite result would be expected. Because PMA also targets eukaryotic DNA, it is probable that there was a substantial number of damaged host cells during the tissue removal via air brushing that were labelled by PMA. This would in turn shift the proportion of bacteria relative to coral cells to be greater in the PMA treatment and result in no difference when looking at the ratio of *Endozoicomonas* to all bacteria. Possible ways to mitigate this issue are to use normalization approaches that are not dependent on DNA, such as surface area of the collected fragment or by the number of coral polyps. While it was not the motivation of this study, one advantage of the host DNA depletion by PMA is the application of shotgun metagenomics. Eukaryotic DNA can dominate coral samples, obscuring changes in microbial populations because few DNA sequence reads are from the microbial component [110]. PMA treatment after osmotic lysis of human saliva samples has been a successful approach to remove host-derived metagenomic sequences [111] and could be an option to deplete coral DNA from coral tissue as well.

In addition to PCR, successful PMA treatment has been reported in combination with loop-mediated isothermal amplification [112], multiple displacement amplification [113], flow cytometry [33, 114], and metagenomics [111, 114, 115]. There are other promising applications such as PMA in combination with fluorescence *in situ* hybridization (PMA-FISH) [116] that can be explored in future work.

## Conclusions

Coral microbiomes are dynamic ecosystems, composed of tens to hundreds of unique microbial taxa. Metabarcoding of 16S rRNA genes has been widely adopted to understand this diversity more generally, making the profiling of existing microbiomes in different host species a common analysis in the coral field [117]. The function of these microbiomes depends on which members of the community are alive or dead. While only the bacterial communities of *A. loripes* and *P. acuta* were significantly changed by PMA, any removal of non-viable signal has the potential to increase knowledge of the potentially metabolically active microbes. These results provide novel insights into the dynamic nature of host-associated microbiomes and underline the importance of applying versatile tools in the analysis of metabarcoding or next-generation sequencing data sets.

## Declarations

### Ethics approval and consent to participate

Not applicable

### Consent for publication

Not applicable

### Availability of data and material

The datasets generated and analysed during the current study are available under NCBI BioProject ID PRJNA971764.

### Authors’ contributions

AMD conceived the project; AMD, LLB, and MJHvO designed the study; AMD, LG, and AW collected and processed samples through PMA treatment and metabarcoding library prep; ECF completed ddPCR; AMD analysed and interpreted the data; AMD wrote the first draft; all authors substantively revised the manuscript and approve of the submitted version.

## Acknowledgements

The corals used in this study were originally collected from the sea estates of the Wuthathi, Bindal, Wulgurukaba, and Manbarra People. We acknowledge their contributions to this research not only by way of the coral samples, but also through Indigenous Knowledge and custodianship of the land and sea country on which we work. We are grateful to Lonidas Koukoumaftsis, Dr. Jose Montalvo-Proano, Dr. Thomas (Ed) Roberts, Rachel Neil, and Hugo Scharfenstein for coral material.

