## Supplemental Figures for "DNA from non-viable bacteria biases diversity estimates in the corals *Acropora loripes* and *Pocillopora acuta*"

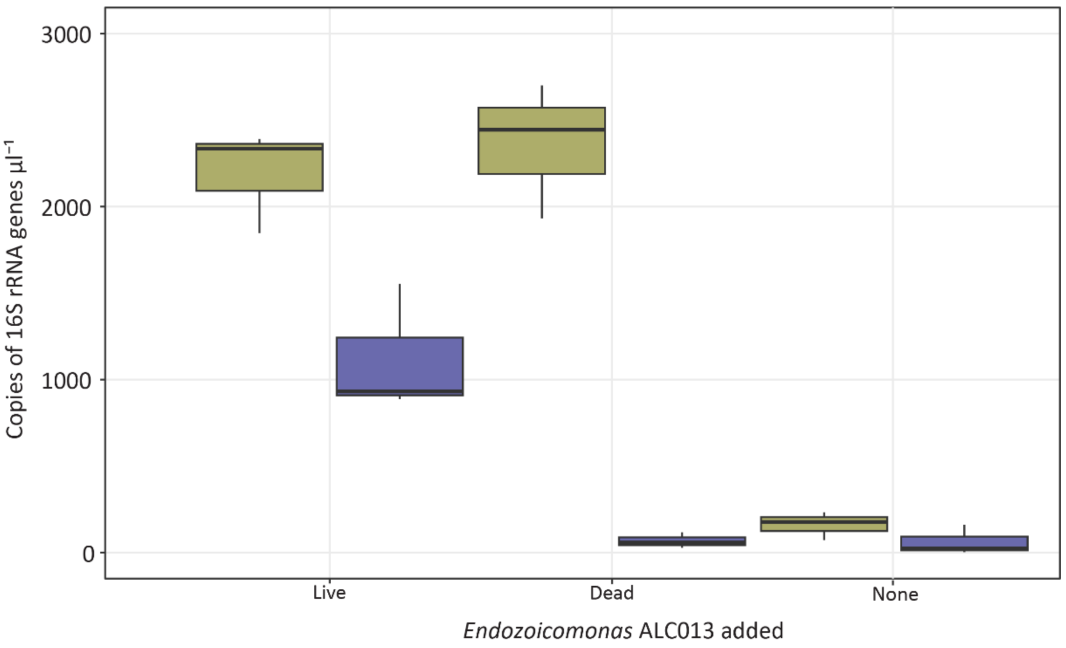


Figure S1: 16S rRNA gene copies $\mu l^{-1}$ of sample from ddPCR for heat-killed anemone homogenate spiked with viable (Live) or heat-killed (Dead) *Endozoicomonas* sp. or unspiked (None) and either untreated (gold) or PMA treated (purple). PMA treatment successfully precluded PCR when samples were spiked with heat-killed bacterial cells.

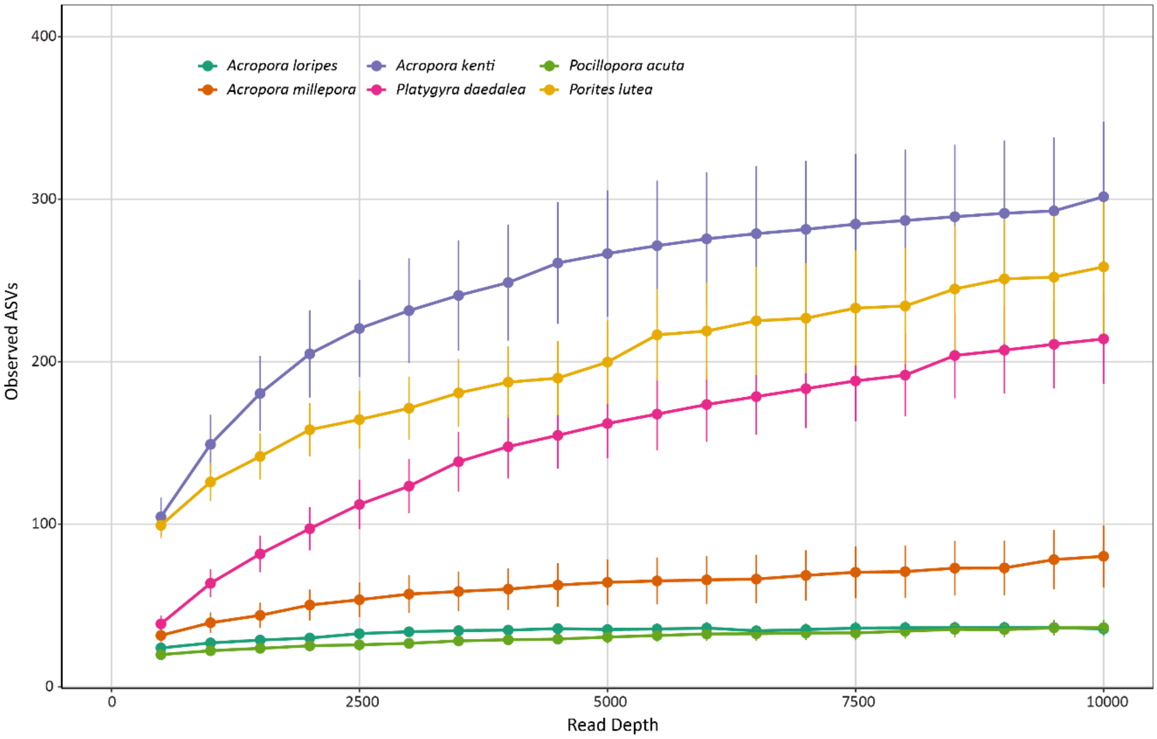


Figure S2: Rarefaction curve showing the relationship between sequencing depth and species richness (Observed ASVs) for each of six coral species sampled (from most to least diverse: *Acropora kenti* – purple; *Porites lutea* – yellow; *Platygyra daedalea* – pink; *A. millepora* – orange; *A. loripes* – dark green*; Pocillopora acuta* – light green). Rarefactions were computed in QIIME2 using the 'alpha-rarefaction' function with ten iterations at intervals of 500 up to 10000 reads sampled. Standard error for each point is shown with the error bars.

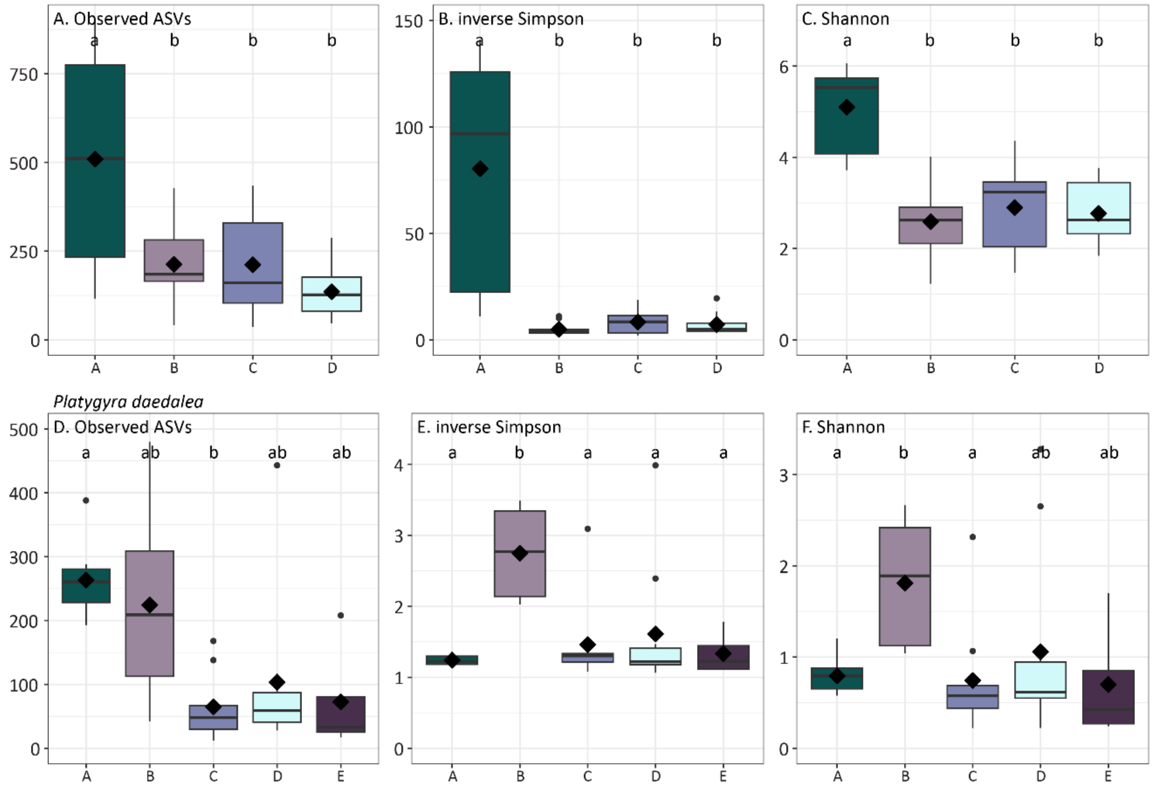


Figure S3: Alpha diversity indices, observed ASVs (A,D), inverse Simpson's index (B,E), and Shannon's index (C,F) by coral genotype for *A. kenti* (A-C), and *P. daedalea* (D-F). Boxes cover the interquartile range (IQR) and the diamond inside the box denotes the median. Whiskers represent the lowest and highest values within 1.5 × IQR. Different small letters indicate significant differences in Tukey's honest significant difference (HSD) post hoc tests.

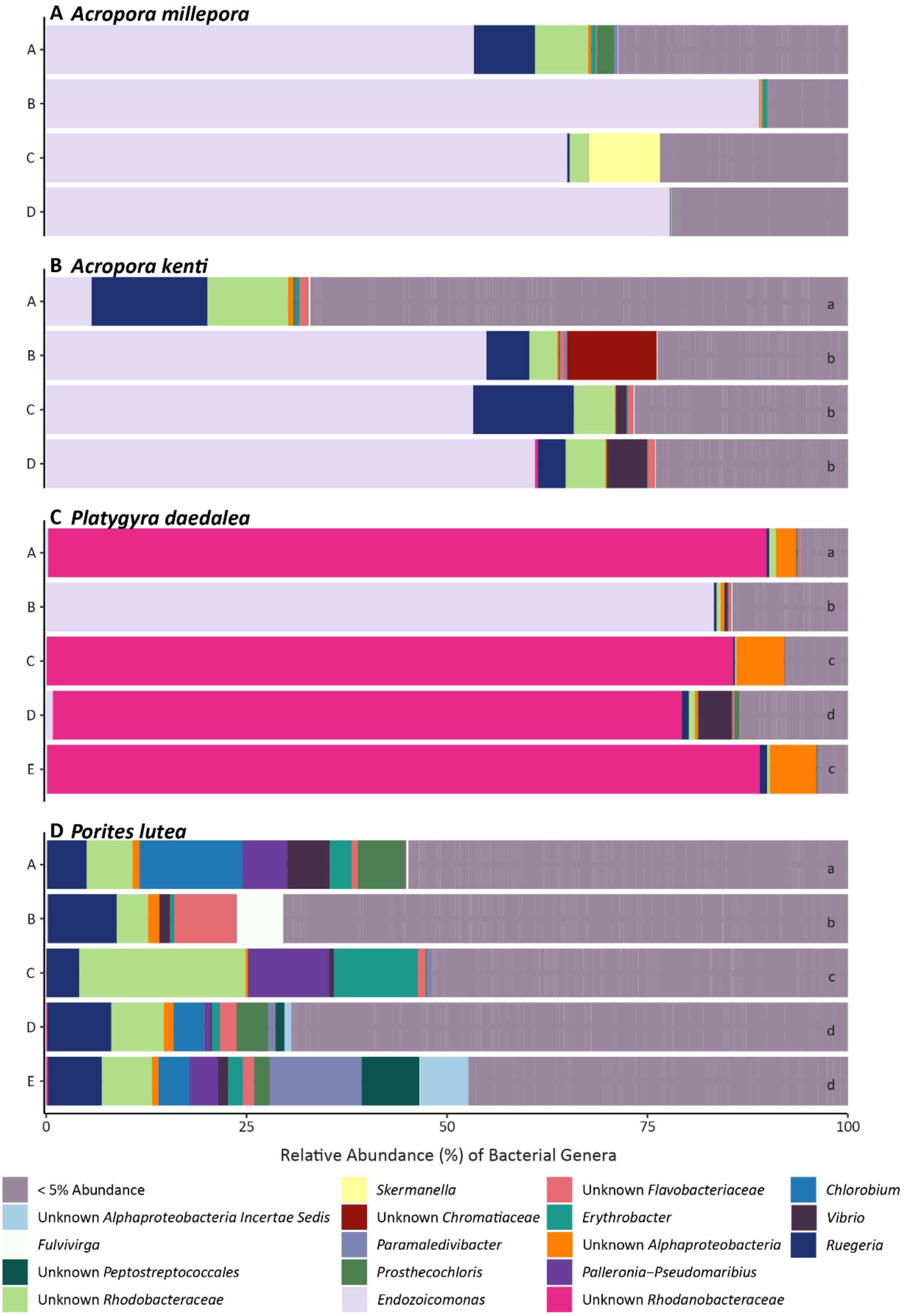


Figure S4: Relative abundance of bacterial genera for *A. millepora* (A), *A. kenti* (B), *P. daedalea* (C), and *P. lutea* (D) by genotype. PMA-treated and untreated samples were pooled as there were no significant differences in community composition for these species. Low abundance genera were pooled into a single category for each species. Where there was a significant difference in bacterial community structure by PERMANOVA, Tukey HSD letters are provided at the right of the barplot. Genotypes within a host species with different lowercase letters indicates that they are significantly different from one another.

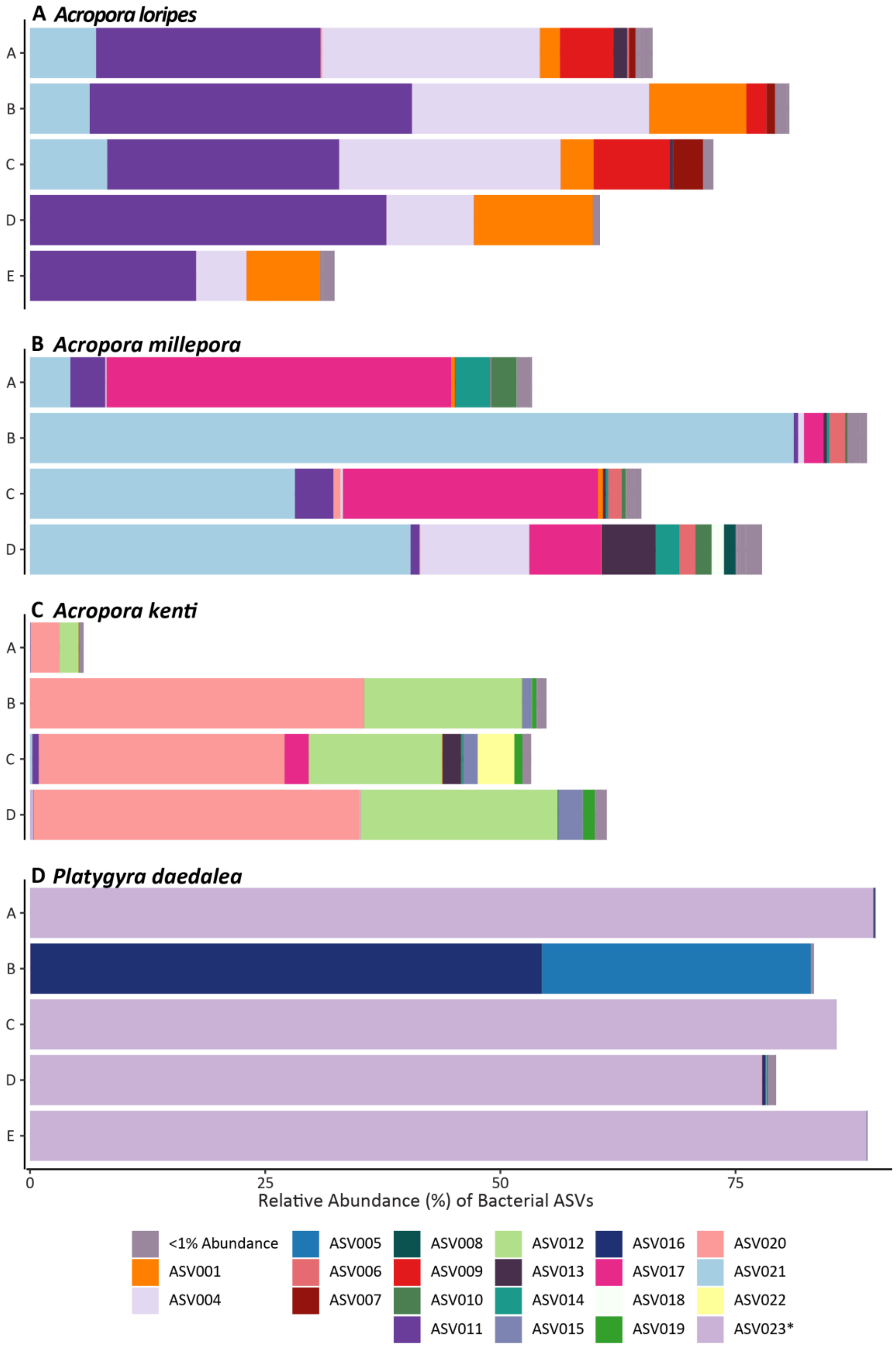


Figure S5: Relative abundance of *Endozoicomonas* (ASV001, ASV004-22) and unknown *Rhodanobacteraceae* (ASV023* only) ASVs in A) *A. loripes*, B) *A. millepora*, C) *A. kenti*, and D)

*P. daedalea* by coral genotype. Untreated and PMA-treated samples were pooled for this visualization.
