## Supplemental Tables for "DNA from non-viable bacteria biases diversity estimates in the corals *Acropora loripes* and *Pocillopora acuta*"

Table S1: Collection and sampling details for all coral colonies used in this study. All corals were collected under permit# G12/35236.1. *After sampling, two colonies of *A. millepora* were identified as belonging to the same genotype.

| Species | Genotype | Date Collected | Reef Site Collected From | Latitude | Longitude | AIMS Holding Tank | Date Sampled | Fragments Sampled (n) |
|---|---|---|---|---|---|---|---|---|
| *Acropora loripes* | A | 20/06/2020 | Davies Reef | 18° 48'S | 147° 39'E | ISB1-A | 17/06/2022 | 5 |
| *Acropora loripes* | B | 20/06/2020 | Davies Reef | 18° 48'S | 147° 39'E | ISB1-A | 17/06/2022 | 5 |
| *Acropora loripes* | C | 5/12/2020 | Backnumbers Reef | 18° 51'S | 147° 15'E | ISB1-A | 17/06/2022 | 5 |
| *Acropora loripes* | D | 5/12/2020 | Backnumbers Reef | 18° 51'S | 147° 15'E | OHP1-SUMP | 17/06/2022 | 5 |
| *Acropora loripes* | E | 20/06/2020 | Davies Reef | 18° 48'S | 147° 39'E | OHP1-SUMP | 17/06/2022 | 5 |
| *Acropora millepora* | A | 1/04/2022 | Davies Reef | 18° 48'S | 147° 39'E | M1S8-A/SGS2-C | 15/06/2022 | 10* |
| *Acropora millepora* | B | 1/04/2022 | Davies Reef | 18° 48'S | 147° 39'E | M1S8-A | 15/06/2022 | 5 |
| *Acropora millepora* | C | 1/04/2022 | Davies Reef | 18° 48'S | 147° 39'E | M1S8-A | 15/06/2022 | 5 |
| *Acropora millepora* | D | 1/04/2022 | Davies Reef | 18° 48'S | 147° 39'E | M1S8-A | 15/06/2022 | 5 |
| *Acropora kenti* | A | 22/02/2022 | Davies Reef | 18° 48'S | 147° 39'E | OHP1-SUMP | 16/06/2022 | 5 |
| *Acropora kenti* | B | 22/02/2022 | Davies Reef | 18° 48'S | 147° 39'E | OHP1-SUMP | 16/06/2022 | 5 |
| *Acropora kenti* | C | 22/02/2022 | Davies Reef | 18° 48'S | 147° 39'E | OHP1-SUMP | 16/06/2022 | 5 |
| *Acropora kenti* | D | 22/02/2022 | Davies Reef | 18° 48'S | 147° 39'E | OHP1-SUMP | 16/06/2022 | 5 |
| *Platygyra daedalea* | A | 1/09/2021 | Davies Reef | 18° 48'S | 147° 39'E | OHP1-SUMP | 16/06/2022 | 5 |
| *Platygyra daedalea* | B | 1/09/2021 | Davies Reef | 18° 48'S | 147° 39'E | OHP1-SUMP | 16/06/2022 | 5 |
| *Platygyra daedalea* | C | 1/09/2021 | Davies Reef | 18° 48'S | 147° 39'E | OHP1-SUMP | 16/06/2022 | 5 |
| *Platygyra daedalea* | D | 1/09/2021 | Davies Reef | 18° 48'S | 147° 39'E | OHP1-SUMP | 16/06/2022 | 5 |
| *Platygyra daedalea* | E | 1/09/2021 | Davies Reef | 18° 48'S | 147° 39'E | OHP1-SUMP | 16/06/2022 | 5 |
| *Pocillopora acuta* | A | 20/07/2017 | Rib Reef | 18° 47'S | 146° 87'E | 1HP1-E | 15/06/2022 | 5 |
| *Pocillopora acuta* | B | 20/07/2017 | Rib Reef | 18° 47'S | 146° 87'E | 1HP1-F | 15/06/2022 | 5 |
| *Pocillopora acuta* | C | 20/07/2017 | Rib Reef | 18° 47'S | 146° 87'E | 1HP1-F | 15/06/2022 | 5 |
| *Pocillopora acuta* | D | 20/07/2017 | Rib Reef | 18° 47'S | 146° 87'E | 1HP1-G | 15/06/2022 | 5 |
| *Pocillopora acuta* | E | 20/07/2017 | Rib Reef | 18° 47'S | 146° 87'E | 1HP1-G | 15/06/2022 | 5 |
| *Porites lutea* | A | 9/11/2021 | Wood Reef | 11° 80'S | 143° 97'E | OHP1-SUMP | 16/06/2022 | 5 |
| *Porites lutea* | B | 9/11/2021 | Wood Reef | 11° 80'S | 143° 97'E | OHP1-SUMP | 16/06/2022 | 5 |

| | | | | | | | | |
|---|---|---|---|---|---|---|---|---|
| *Porites lutea* | C | 9/11/2021 | Wood Reef | 11° 80'S | 143° 97'E | OHP1-SUMP | 16/06/2022 | 5 |
| *Porites lutea* | D | 9/11/2021 | Wood Reef | 11° 80'S | 143° 97'E | OHP1-SUMP | 16/06/2022 | 5 |
| *Porites lutea* | E | 9/11/2021 | Wood Reef | 11° 80'S | 143° 97'E | OHP1-SUMP | 16/06/2022 | 5 |

Table S2: Putative contaminants identified with decontam by source (PCR, DNA extraction, or tissue blasting) with their relative abundance in the entire dataset before removal.

| ASV | Relative Abundance (%) | Source | Phylum | Family | Genus |
|---|---|---|---|---|---|
| Contam001 | 0.003 | PCRNeg | *Actinobacteriota* | *Corynebacteriaceae* | *Corynebacterium* |
| Contam002 | 0.001 | PCRNeg | *Planctomycetota* | *Pirellulaceae* | *Pir4_lineage* |
| Contam003 | 0.004 | PCRNeg | *Myxococcota* | bacteriap25 | bacteriap25 |
| Contam004 | 0.007 | PCRNeg | *Bacteroidota* | *Flavobacteriaceae* | Unknown *Flavobacteriaceae* |
| Contam005 | 0.001 | PCRNeg | *Proteobacteria* | *Legionellaceae* | *Legionella* |
| Contam006 | 0.001 | PCRNeg | *Proteobacteria* | *Geminicoccaceae* | Unknown *Geminicoccaceae* |
| Contam007 | 0.003 | PCRNeg | *Proteobacteria* | *Rhodobacteraceae* | *Limimaricola* |
| Contam008 | 0.001 | PCRNeg | *Proteobacteria* | *Kiloniellaceae* | Unknown *Kiloniellaceae* |
| Contam009 | 0.009 | Ext_Blank | *Actinobacteriota* | *Micrococcaceae* | *Kocuria* |
| Contam010 | 0.015 | Ext_Blank | *Actinobacteriota* | *Actinomycetaceae* | *Actinomyces* |
| Contam011 | 0.003 | Ext_Blank | *Firmicutes* | *Streptococcaceae* | *Streptococcus* |
| Contam012 | 0.008 | Ext_Blank | *Bacteroidota* | *Amoebophilaceae* | *Candidatus* Amoebophilus |
| Contam013 | 0.003 | Ext_Blank | *Proteobacteria* | *Pseudomonadaceae* | *Pseudomonas* |
| Contam014 | 0.017 | Ext_Blank | *Proteobacteria* | *Comamonadaceae* | *Schlegelella* |
| Contam015 | 0.006 | Ext_Blank | *Proteobacteria* | *Xanthobacteraceae* | *Afipia* |
| Contam016 | 0.006 | Ext_Blank | *Proteobacteria* | *Xanthobacteraceae* | *Bradyrhizobium* |
| Contam017 | 0.114 | Blank | *Actinobacteriota* | *Micrococcaceae* | *Arthrobacter* |
| Contam018 | 0.051 | Blank | *Actinobacteriota* | *Nocardioidaceae* | *Nocardioides* |
| Contam019 | 0.000 | Blank | *Verrucomicrobiota* | cvE6 | cvE6 |
| Contam020 | 0.004 | Blank | *Dependentiae* | *Vermiphilaceae* | *Vermiphilaceae* |
| Contam021 | 0.006 | Blank | *Firmicutes* | *Lachnospiraceae* | *Epulopiscium* |
| Contam022 | 0.001 | Blank | *Bacteroidota* | *Amoebophilaceae* | Unknown *Amoebophilaceae* |

| | | | | | |
|---|---|---|---|---|---|
| Contam023 | 0.000 | Blank | *Proteobacteria* | *Acetobacteraceae* | *Acidiphilium* |
| Contam024 | 0.001 | Blank | *Proteobacteria* | *Beijerinckiaceae* | *Methylobacterium-Methylorubrum* |
| Contam025 | 0.002 | Blank | *Proteobacteria* | *Rhodobacteraceae* | *Amaricoccus* |
| Contam026 | 0.058 | Blank | *Proteobacteria* | *Rhodobacteraceae* | *Amaricoccus* |
| Contam027 | 0.268 | Blank | *Proteobacteria* | *Azospirillaceae* | *Skermanella* |
| **Total Contamination (%)** | **0.592** | | | | |

Table S3: β-diversity pairwise comparisons by genotype for *A. kenti*, *P. daedalea*, and *P. lutea*, which were completed using a weighted unifrac distance matrix and the package pairwise.adonis. p values have been corrected based on a Holm adjustment method. Tukey HSD letters were assigned to each genotype based on these statistics and added to barplots in Fig. S4.

| Species | Genotype Pairs | SumsofSqs | F model | R2 | Holm p value |
|---|---|---|---|---|---|
| *A. kenti* | A vs B | 0.113 | 20.807 | 0.536 | 0.0006 |
| *A. kenti* | A vs C | 0.102 | 17.212 | 0.489 | 0.0006 |
| *A. kenti* | A vs D | 0.116 | 28.025 | 0.609 | 0.0006 |
| *A. kenti* | B vs C | 0.007 | 1.069 | 0.056 | 0.6316 |
| *A. kenti* | B vs D | 0.011 | 2.298 | 0.113 | 0.1911 |
| *A. kenti* | C vs D | 0.006 | 1.084 | 0.057 | 0.6316 |
| *P. daedalea* | A vs B | 0.181 | 599.313 | 0.971 | 0.0010 |
| *P. daedalea* | A vs C | 0.000 | 2.061 | 0.103 | 0.0384 |
| *P. daedalea* | A vs D | 0.000 | 5.172 | 0.223 | 0.0012 |
| *P. daedalea* | A vs E | 0.000 | 3.446 | 0.198 | 0.0060 |
| *P. daedalea* | B vs C | 0.182 | 554.484 | 0.969 | 0.0010 |
| *P. daedalea* | B vs D | 0.180 | 525.621 | 0.967 | 0.0010 |
| *P. daedalea* | B vs E | 0.135 | 342.766 | 0.961 | 0.0010 |
| *P. daedalea* | C vs D | 0.000 | 3.953 | 0.180 | 0.0050 |
| *P. daedalea* | C vs E | 0.000 | 0.676 | 0.046 | 0.6800 |
| *P. daedalea* | D vs E | 0.000 | 3.501 | 0.200 | 0.0384 |

| | | | | | |
|---|---|---|---|---|---|
| *P. lutea* | A vs B | 0.008 | 2.154 | 0.107 | 0.0136 |
| *P. lutea* | A vs C | 0.010 | 3.566 | 0.165 | 0.0014 |
| *P. lutea* | A vs D | 0.008 | 2.173 | 0.108 | 0.0261 |
| *P. lutea* | A vs E | 0.013 | 4.540 | 0.201 | 0.0014 |
| *P. lutea* | B vs C | 0.008 | 3.026 | 0.144 | 0.0014 |
| *P. lutea* | B vs D | 0.009 | 2.411 | 0.118 | 0.0010 |
| *P. lutea* | B vs E | 0.010 | 3.994 | 0.182 | 0.0010 |
| *P. lutea* | C vs D | 0.005 | 1.795 | 0.091 | 0.0261 |
| *P. lutea* | C vs E | 0.007 | 4.371 | 0.195 | 0.0010 |
| *P. lutea* | D vs E | 0.004 | 1.580 | 0.081 | 0.0510 |

Table S4: Review of studies examining non-viable versus viable bacteria by the use of PMA in mixed communities.

| Community Evaluated | [PMA], Incubation, and Light Conditions | Main finding | Quantification Method | Reference |
|---|---|---|---|---|
| Coral (*Acropora loripes*, *A. millepora*, *A. kenti*, *Platygyra daedalea*, *Pocillopora acuta*, and *Porites lutea*) | 25 µM PMAxx in the dark at RT for 10 min with constant agitation; 30 min light in PMA-Lite™ LED photolysis device, mixing every 3 min | DNA from non-viable microbes significantly inflated community evenness (85%) and species diversity (31%) for *A. loripes* with significant differences in bacterial community structure between PMA-treated and-untreated samples in *A. loripes* and *P. acuta*. | ddPCR | This study |
| Fresh equine manure | 80 µM PMAxx in the dark at RT for 10 min vortexing for several seconds every 2 min; 15 min light in PMA-Lite™ LED photolysis device, rotating every 3 min | No significant differences in ecological indices or mean estimated total living bacteria were found in the final faecal filtrate compared to the original manure. | qPCR | [1] |
| Rainwater | 50 µM PMAxx in the dark at RT for 10 min; 5 min exposure to 500 W light on ice ~20 cm from the light source | Metabarcoding β-diversity indices indicated that, in comparison to the untreated samples, PMA reduced the detection of nonviable bacteria in the rainwater samples. | qPCR | [2, 3] |
| Human saliva | 10 µM PMAxx in the dark at RT for 5 min; 25 min exposure to benchtop fluorescent light bulb with tubes horizontally on ice at a | Removing relic DNA from saliva samples did not greatly impact the microbial composition, it did | Flow cytometry and qPCR | [4] |

| | | | | |
|---|---|---|---|---|
| | distance <20 cm from the light source with a brief mixing every 5-10 min | increase our resolution among samples collected over time. | | |
| Farmed yellowtail kingfish (*Seriola lalandi*) gut and mucosa | 10 µM PMAxx in the dark at RT for 5 min; 25 min exposure to two 500 W halogen globes with tubes horizontally on ice at a distance <30 cm from the light source with a brief mixing every 5 min | PMA treatment significantly reduced the microbial diversity and richness of digesta and mucosal samples and depleted bacterial constituents typically considered to be important within fish, such as *Lactobacillales* and *Clostridales*. | NA | [5] |
| Juvenile Atlantic Salmon gut | 50 µM PMAxx in the dark at RT for 5 min; "exposure to light for 30 min in a lightbox from Geniul" | 9.1% of the sequencing reads came from nonviable bacterial cells. | NA | [6] |
| Faeces from 16 human subjects | 50 µM PMAxx in the dark at 4 °C for 10 min; 10 min light in PMA-Lite™ LED photolysis device "this procedure was repeated 3 times…" | 40% of DNA in faecal samples can be attributed to extracellular DNA and damaged bacterial cells | Flow cytometry and qPCR | [7] |
| Anaerobic digestor sludge | 100 µM PMAxx at RT for 5 min; 15 min light in PMA-Lite™ LED photolysis device, mixing every 5 min | Quantitative PCR revealed that 5–30% of the rRNA genes were derived from inactive or dead cells in anaerobic sludge digesters. | qPCR | [8] |
| Meconium | 50 µM PMAxx in the dark at 37 °C for 15 min with occasional vortexing; 15 min light in PMA-Lite™ LED photolysis device, mixing every 5 min | PMA-treated meconium samples did not differ significantly from untreated samples in terms of observed number of OTUs; although they did differ taxonomically, with around one quarter of OTUs identified in untreated samples only, suggesting that they have originated from cell-free/nonviable DNA. | qPCR | [9] |
| Gut of Rex rabbits | 100 µM PMAxx in the dark at RT for 5 min with occasional inversion; samples were then transferred into transparent bags and flattened to an approximately 0.1 mm thick layer. For PMA activation, the samples were exposed to a 650 W halogen light source for 8 min on ice 20 cm below the light source. | The alpha- and beta-diversities differed significantly between groups. Many dead bacteria existed in the digestive tract of Rex rabbits and distorted the community profile of the live microbiota. Total bacteria are an improper representation of the live gut microbiota, particularly in the foregut. | qPCR | [10] |

| | | | | |
|---|---|---|---|---|
| Shrimp – food spoilage | 100 µM PMAxx in the dark at RT for 10 min with constant agitation; 15 min light in PMA-Lite™ LED photolysis device with occasional shaking | Shannon-Wiener index and PCoA analysis indicated there were significant differences between bacterial diversity in samples treated with and without PMA. | NA | [11] |
| Soil | 40 µM PMAxx in the dark at RT for 4 min with constant gentle vortexing. For PMA activation, the samples were exposed to a 650 W halogen light source for four consecutive 30 s/30 s light/dark cycles, while continually vortexing, 20 cm below the light source. | 40% of both prokaryotic and fungal DNA was extracellular or from cells that were no longer intact. Extracellular DNA inflated the observed prokaryotic and fungal richness by up to 55% and caused significant misestimation of taxon relative abundances, including the relative abundances of taxa integral to key ecosystem processes. | qPCR | [12] |

Supplemental Tables References:


1. Loublier C, Taminiau B, Heinen J, Lecoq L, Amory H, Daube G, Cesarini C: **Evaluation of Bacterial Composition and Viability of Equine Feces after Processing for Transplantation.** *Microorganisms* 2023, **11**.
2. Reyneke B, Ndlovu T, Khan S, Khan W: **Comparison of EMA-, PMA- and DNase qPCR for the determination of microbial cell viability.** *Appl Microbiol Biotechnol* 2017, **101:**7371-7383.
3. Reyneke B, Waso M, Ndlovu T, Clements T, Havenga B, Khan S, Khan W: **EMA- Versus PMA-Amplicon-Based Sequencing to Elucidate the Viable Bacterial Community in Rainwater.** *Water, Air, & Soil Pollution* 2022, **233**.
4. Marotz C, Morton JT, Navarro P, Coker J, Belda-Ferre P, Knight R, Zengler K: **Quantifying Live Microbial Load in Human Saliva Samples over Time Reveals Stable Composition and Dynamic Load.** *mSystems* 2021, **6**.
5. Legrand T, Wos-Oxley ML, Wynne JW, Weyrich LS, Oxley APA: **Dead or alive: microbial viability treatment reveals both active and inactive bacterial constituents in the fish gut microbiota.** *J Appl Microbiol* 2021.
6. Dvergedal H, Sandve SR, Angell IL, Klemetsdal G, Rudi K: **Association of gut microbiota with metabolism in juvenile Atlantic salmon.** *Microbiome* 2020, **8:**160.
7. Galazzo G, van Best N, Benedikter BJ, Janssen K, Bervoets L, Driessen C, Oomen M, Lucchesi M, van Eijck PH, Becker HEF, et al: **How to Count Our Microbes? The Effect of Different Quantitative Microbiome Profiling Approaches.** *Front Cell Infect Microbiol* 2020, **10:**403.
8. Ni J, Hatori S, Wang Y, Li YY, Kubota K: **Uncovering Viable Microbiome in Anaerobic Sludge Digesters by Propidium Monoazide (PMA)-PCR.** *Microb Ecol* 2020, **79:**925-932.
9. Stinson LF, Keelan JA, Payne MS: **Characterization of the bacterial microbiome in first-pass meconium using propidium monoazide (PMA) to exclude nonviable bacterial DNA.** *Lett Appl Microbiol* 2019, **68:**378-385.
10. Fu X, Zeng B, Wang P, Wang L, Wen B, Li Y, Liu H, Bai S, Jia G: **Microbiome of Total Versus Live Bacteria in the Gut of Rex Rabbits.** *Front Microbiol* 2018, **9:**733.
11. Zhao F, Liu H, Zhang Z, Xiao L, Sun X, Xie J, Pan Y, Zhao Y: **Reducing bias in complex microbial community analysis in shrimp based on propidium monoazide combined with PCR-DGGE.** *Food Control* 2016, **68:**139-144.
12. Carini P, Marsden PJ, Leff JW, Morgan EE, Strickland MS, Fierer N: **Relic DNA is abundant in soil and obscures estimates of soil microbial diversity.** *Nat Microbiol* 2016, **2:**16242.
